# The CYP1A1 connection between enterolactone and breast cancer risk

**DOI:** 10.1101/2024.06.06.597836

**Authors:** Juana Hatwik, Ningthoujam Sonia, Anil M. Limaye

**Affiliations:** Department of Biosciences and Bioengineering, Indian Institute of Technology Guwahati, Guwahati 781039, Assam, India; Department of Health Sciences, Al-Baath University, Homs, Syria

**Keywords:** enterolactone, lignans, breast cancer, CYP1A1, AHR

## Abstract

Enterolactone (EL), a mammalian enterolignan, is a gut microbe-generated plant-lignan derivative. Although observational data are controversial, meta-analyses have affirmed that plant lignan-rich diets, or high serum EL reduce breast cancer risk, or the associated mortality, in post-menopausal women. However, the mechanistic basis is unknown. Here we show that EL antagonizes AHR to reduce *CYP1A1* mRNA expression in MCF-7 breast cancer cells. Intriguingly, it increases CYP1A1 protein, a mediator of xenobiotic response, which is frequently expressed in breast tumors, and implicated in cell proliferation and survival. EL’s effect on CYP1A1 expression is similar to estrogen, and mediated via ERα. But, by virtue of partial ERα agonism/antagonism, EL attenuates estrogen-mediated increase in CYP1A1 protein. These data suggest potential mechanisms underlying EL’s beneficial effects in breast cancer. In the face of xenobiotic exposure, its AHR antagonism may reduce the generation of cancer-inducing genotoxic agents. With declining levels of estrogen in post-menopausal women, EL may antagonize estrogen-mediated induction of CYP1A1 protein, and the associated proliferation of mammary epithelial cells.

## Introduction

Meta-analyses of the observational data have revealed that lignan intake, or high serum enterolactone (EL), reduces the risk of breast cancer, in post-menopausal women (1). EL is generated by the gut microbes through biotransformation of dietary plant lignans. Breast cancer risk-reduction is generally attributed to the anti-proliferative action of EL (2). However, its estrogen-like properties, including enhanced proliferation of ERα-positive breast cancer cells (3, 4) appear counterintuitive. The basis of cancer risk-reducing potential of EL remains unknown.

## Results and Discussion

MCF-7 cells treated with 10 µM EL are characterized by negative enrichment of hallmark xenobiotic metabolism genes compared to control (4). The leading edge genes of this gene-set contained few cytochrome P450 family members, including CYP1A1, and other xenobiotic response genes, which are downmodulated by EL (SI Appendix II, Fig. S1). 17β-estradiol (E2) downmodulates *CYP1A1* mRNA (5). In keeping with its known estrogenicity (3, 4), EL, like E2 (SI Appendix II, Fig. S2), downmodulated the expression of *CYP1A1* mRNA, but increased CYP1A1 protein (Fig. 1A,B). Tamoxifen, a selective estrogen receptor modulator significantly blocked EL-mediated decrease in *CYP1A1* mRNA (SI Appendix II, Fig. S3). Fulvestrant, a selective estrogen receptor degrader, also blocked EL-mediated modulation of *CYP1A1* mRNA or protein (Fig. 1C). These observations strongly supported the role of ERα. Analysis of ChIP-seq data (GSE25710) revealed two sites of estrogen-induced ERα occupancy in the *CYP1A1* locus (site 1 and site 2, Fig. 1D). ChIP experiments employing primers targeting site 1 (Fig. 1D), which contained an estrogen response element (ERE) predicted by JASPAR tool, confirmed that EL-mediated downmodulation was associated with enhanced ERα binding (Fig. 1E, left panel).

**Fig. 1.**
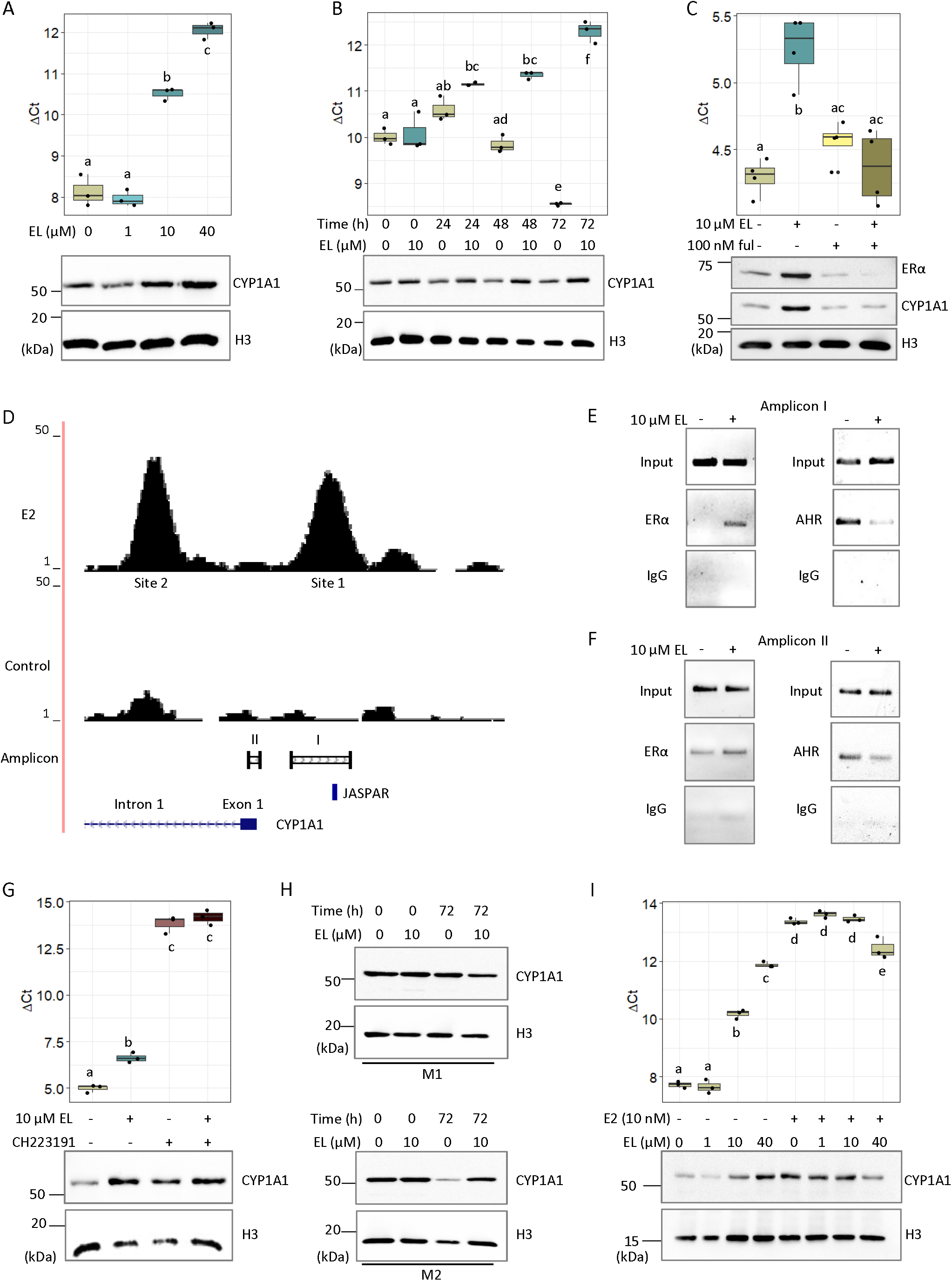
EL differentially modulates CYP1A1 mRNA and protein via ERα in MCF-7 breast cancer cells. **A, B**. Results of dose-response (**A**) and time-course experiments (**B**) to study the effect of EL on *CYP1A1* mRNA (top) and protein expression (bottom). **C**. Fulvestrant (ful) pre-treatment significantly blocks the effect of EL on *CYP1A1* mRNA (top) and protein expression (bottom). **D**. A snapshot of the *CYP1A1* locus. Site 1, and Site 2 are ERα enriched regions deduced from the ChIP-seq data (GSE25710). ERE predicted by JASPAR is indicated in blue. The amplicon track shows the regions amplified by two pairs of ChIP primers (labeled as I and II). **E, F**. Results of ChIP experiments with primer pairs generating amplicon I (**E**), or amplicon II (**F**), respectively. **G**. Effect of CH223191 on EL’s modulation of *CYP1A1* mRNA (top) and protein (bottom). **H**. EL restores CYP1A1 protein expression in cells grown in M2. Cells were treated with 0.1% DMSO (vehicle) or 10 μM EL for 0 or 72 h, in M1 or M2 medium. **I**. EL reverses the effect of E2 on CYP1A1 mRNA (top) and protein (bottom).

Compelling experimental data show that the interplay of opposing estrogenic and xenobiotic signals determines the net *CYP1A1* mRNA expression, wherein, estrogen-activated ERα directly interferes with xenobiotic-induced, and AHR-dependent transcriptional activation of *CYP1A1* (6). Given that experiments discussed here were conducted without xenobiotic exposure, both EL, and E2 impacted the basal CYP1A1 expression. A precipitous fall in *CYP1A1* mRNA levels by CH223191, an inhibitor of AHR nuclear translocation (Fig. 1G), showed that basal *CYP1A1* mRNA expression is AHR-dependent. EL-treatment, over and above CH223191, did not reduce it any further. We hypothesized that EL-mediated ERα activation interferes with AHR-mediated transactivation at the *CYP1A1* locus. Concomitant increase in ERα, and decrease in AHR occupancy, at site 1 (Fig. 1E) or the site of interference demonstrated by Marques et al (6) (Fig. 1F), strongly supported the hypothesis.

Overall, EL appeared to mimic estrogen action at the *CYP1A1* locus; except that EL increased ERα protein (Fig. 1C), while E2 is known to decrease it (7). How both increase or decrease of ERα protein could be associated with altered CYP1A1 expression, is not clear. However, EL-mediated increase in ERα protein, on its own, may suppress *CYP1A1* mRNA, because, a mere reduction in ERα protein (with fulvestrant) significantly increased AHR protein, and *CYP1A1* mRNA (SI Appendix II, Fig. S4). CYP1A1 is a part of xenobiotic response. It metabolizes pro-carcinogens into carcinogens (8). However, it is one among a large repertoire of metabolic enzymes. Besides CYP1A1, other cytochrome P450 family members, such as CYP2S1, CYP27A1, CYP26A1, and CYP2E1, are also a part of the leading edge genes within the negatively enriched xenobiotic response gene-set (SI Appendix II, Fig. S1). Since these enzymes acting on endogenous and exogenous substrates can potentially lead to generation of reactive intermediates (9), downmodulation of these genes possibly explain, at least in part, EL’s beneficial effects in breast cancer. But, the induction of CYP1A1 protein by EL obscures the relevance of CYP1A1 in the context of EL’s protective effect. Firstly, enhanced CYP1A1 protein can lead to enhanced production of carcinogens. Secondly, CYP1A1 converts E2 to 2-hydroxy-estradiol (2-OHE2), which is more reactive and may promote carcinogenesis by facilitating the generation of reactive oxygen species. On the other hand, domination of CYP1A1 activity over CYP1B1, which converts E2 to 4-hydroxy-estradiol (4-OHE2), is beneficial from the cancer perspective; given the anti-proliferative and genotoxic nature of the 2-OHE2 and 4-OHE2, respectively (10). Thus, any explanation for the beneficial effects of EL on the basis of *CYP1A1* mRNA or protein modulation may not be straightforward. However, a non-enzymatic activity of CYP1A1 associated to cell survival and proliferation is discussed hereunder, which, in the light of more observations from this study, may be the basis of the protection imparted by EL.

CYP1A1 is expressed in majority of breast tumors (11, 12). Experimental depletion of CYP1A1 expression reduces proliferation and increases apoptosis via AMPK activation, and de-phosphorylation of AKT, ERK1/2 and p70S6K (13). Carnosol, a pharmacological agent that brings down CYP1A1 protein levels, also impairs proliferation (13). The proliferative potential of MCF-7 cells in phenol red-free, and charcoal-stripped serum containing medium (M2) is significantly lower than those grown in routine medium (M1) (4). We have observed that MCF-7 cells grown for 72 h in M2 show less CYP1A1 protein expression than those grown in M1 (Fig. 1H). Moreover, 10 µM EL, which restored growth and PCNA expression in cells grown in M2 (4) also restored CYP1A1 protein (Fig. 1H). E2, which induces the proliferation of ERα-positive breast cancer cells, also increased CYP1A1 protein, although, like EL, it downmodulated the mRNA (SI Appendix II, Fig. S2). The correlation of CYP1A1 protein expression with estrogenic input is also borne out by the observation that breast tumors in pre-menopausal women have higher expression of CYP1A1 protein than those in post-menopausal women (12). Rodriguez et al (13) showed that CYP1A1 protein knockdown, but not inhibition of its enzyme activity, impairs cell proliferation. Thus, CYP1A1 protein is associated with proliferation and survival pathways. Furthermore, cell proliferation and xenobiotic metabolism are mutually exclusive functional attributes of CYP1A1. Given that both estrogen and EL induced CYP1A1 protein, the question as to how EL may exert a protective effect is a question that begs attention. EL is known to exert partial agonist/antagonistic activity (14). In agreement with this notion, we found that although E2 or EL induced the expression of the classical estrogen-regulated *TFF1* mRNA, EL attenuated the induction brought about by E2 (SI Appendix II, Fig. S5). Similarly, while E2 or EL independently decreased or increased the levels of *CYP1A1* mRNA and protein respectively, EL attenuated the E2 effect (Fig. 1I). This parallels the observation that both induce proliferation, but EL can attenuate E2-mediated proliferation (15). This property of EL may be a significant contributor to its beneficial effects. We propose that in the face of diminishing levels of estrogen in post-menopausal women, EL concentrations achieved through lignan-rich diet are better able to antagonize estrogen to keep levels of CYP1A1 protein low, thereby reducing the chances of breast tumors.

In summary, EL acts via ERα to antagonize AHR-mediated transactivation that results in downmodulation of *CYP1A1* mRNA. It possibly imparts beneficial effects by inhibiting the oncogenic effects of genotoxic agents produced by xenobiotic response to pro-carcinogens, such as polycyclic or heterocyclic aromatic hydrocarbons. EL, like E2 induces CYP1A1 protein. However, consistent with the property of partial agonism/antagonism, it blocks E2-mediated increase of CYP1A1 protein, thereby negatively influencing the proliferation of mammary epithelial cells. Therefore, EL may reduce breast cancer risk, irrespective of the menopausal status. However, with diminishing levels of estrogen, EL concentrations achieved with plant lignan-rich diet are better able to antagonize E2 action, that is reflected in significantly reduced breast cancer risk or disease progression in post-menopausal women.

## Materials and Methods

All the experiments were performed on MCF-7 cells. Detailed methodologies are presented as supporting information (SI Appendix I).

## Acknowledgments

The work was supported by financial assistance from Science and Engineering Research Board, Department of Science and Technology, Govt. of India (File No. CRG/2020/002109). Infrastructural support from Department of Biosciences and Bioengineering, IIT Guwahati is acknowledged. JH acknowledges the research fellowship from Al-Baath University, Ministry of Higher Education and Scientific Research, Syrian Arab Republic.

