## Supplementary material for "The CYP1A1 connection between enterolactone and breast cancer risk": SI Appendix I

### SI Appendix I: Materials and Methods

#### Plasticware, chemicals, and reagents

Details of the plasticware, chemicals, and reagents used in this research are provided in **Table S1.**

**Table S1.** List of chemicals and reagents with their sources.

| Item | Cat. No. | Company | Source |
|---|---|---|---|
| DMEM with phenol red | AT-007 | HiMedia | Mumbai, India |
| RPMI-1640 with phenol red | AT-028 | HiMedia | Mumbai, India |
| DMEM without phenol red | AT-190 | HiMedia | Mumbai, India |
| RPMI-1640 without phenol red | AT-120 | HiMedia | Mumbai, India |
| FBS | RM10432 | HiMedia | Mumbai, India |
| CS-FBS | RM10416 | HiMedia | Mumbai, India |
| Trypsin-EDTA | TCL-034 | HiMedia | Mumbai, India |
| Antibiotic solution 100 X liquid (10,000 U penicillin and 10 mg streptomycin per ml in citrate buffer) | A018 | HiMedia | Mumbai, India |
| DPBS | TS-1006 | HiMedia | Mumbai, India |
| Enterolactone | 45199 | Sigma-Aldrich | MO, USA |
| 17β-estradiol | E8875 | Sigma-Aldrich | MO, USA |
| 4-hydroxy-tamoxifen | H7904 | Sigma-Aldrich | MO, USA |
| Fulvesrant | 14409 | Sigma-Aldrich | MO, USA |
| CH223191 | C8124 | Sigma-Aldrich | MO, USA |
| High-Capacity cDNA Reverse transcriptase kit | 4368814 | Applied Biosystems | USA |
| Power Up ™ SYBR Green PCR mix | A25743 | Thermo Scientific | PA, USA |
| Protein G plus-Agarose beads | IP04-1.5ML | Merck Millipore | Burlington, USA |
| Nitrocellulose membrane | SF110B | HiMedia | Mumbai, India |
| ERα antibody (ChIP) | D8H8 | Cell Signaling Technology | MA, USA |
| ERα antibody (Western blot) | 8002 | Santa Cruz Biotechnology | USA |
| AHR antibody | 83200S | Cell Signaling Technology | MA, USA |
| CYP1A1 antibody | PA1-340 | Thermo Scientific | PA, USA |
| Anti-H3 antibody | BB-AB0055 | BioBharti LifeSciences | Kolkata, India |
| Goat anti-rabbit HRP-tagged secondary antibody | 7074S | Cell Signaling Technology | MA, USA |
| Horse anti-mouse HRP-tagged secondary antibody | 7076S | Cell Signaling Technology | MA, USA |
| Clarity Western ECL Substrate | 1705060 | BIO RAD | California, USA |
| Cell culture plasticware | | Eppendorf | Hamburg, Germany |
| | | Thermo Fisher Scientific | PA, USA |
| All other reagents, salts, and buffers | | Merck | Darmstadt, Germany |
| | | Sisco Research Laboratories | Mumbai, India |
| | | Sigma-Aldrich | MO, USA |

**Cell culture**

The ERα-positive MCF-7 breast cancer cells were used in this study. They were grown in a humidified incubator maintained at 37 °C and 5% $CO_2$. Dulbecco Modified Eagle Medium (DMEM), supplemented with 10% fetal bovine serum (FBS), 100 units/mL penicillin, and 100 µg/mL streptomycin (referred to as M1), was used for growth and expansion of cells. For treatments, phenol red-free DMEM supplemented with 10% charcoal-stripped fetal bovine serum (CS-FBS), 100 units/mL penicillin, and 100 µg/mL streptomycin (referred to as M2) was used.

**Experimental protocols**

- **Dose-response study**

  $2 \times 10^5$ cells were seeded in 35 mm dishes using M1. After 50% confluent, cells were washed with DPBS and fed with M2 for 24 h. Thereafter, the cells were treated with 0.1% DMSO (vehicle control), or indicated concentrations of 17β-estradiol (E2) for 24 h, or EL for 72 h, respectively; with the treatment medium being replenished ever 24 h.

- **Time-course study**

  $2 \times 10^5$ cells were seeded in 35 mm dishes using M1, and grown up to 50% confluency. The cells were washed with DPBS and fed with M2 for 24 h. The cells were then treated with 0.1% DMSO (vehicle), or 10 nM E2, or 10 µM EL in M2 for the indicated period. Cells received fresh treatment medium every 24 h.

- **Effect of 4-hydroxy-tamoxifen (4OHT) on EL-mediated suppression of CYP1A1 mRNA**

  Cells were grown upto the desired confluency as above. The cells were then maintained in M2 for 24 h. Thereafter, the cells were treated with 0.1% DMSO (vehicle), 10 µM EL, 1 µM 4OHT, or both using M2 for 72 h, with the treatment medium being changed every 24 h.

- **Effect of CH223191 treatment on EL-mediated modulation of CYP1A1**

  The experiment was conducted as described above for studying the effect of 4-hydroxy-tamoxifen, except that 10 µM CH223191 was used.

- **Effect of fulvestrant treatment on EL-mediated modulation of CYP1A1**

  MCF-7 cells were allowed to grow until 50% confluent. Then they were fed with M2 for 24 h, followed by incubation in M2 with or without 100 nM fulvestrant for another 24 h. After fulvestrant pre-treatment, cells were treated with 0.1% DMSO (vehicle), or 10 µM EL in M2 for 72 h. Cells were fed with fresh M2 supplemented with EL every 24 h.

**RNA extraction, cDNA synthesis, and qPCR**

Total RNA was extracted using a reagent prepared in-house based on Chomczynski and Sacchi (1). 2 µg of total RNA was reverse transcribed using High-Capacity Reverse Transcription kit, as per the manufacturer's instructions. The cDNA was diluted 10 times, and 2 µl (equivalent to 20 ng of total RNA) was used as qPCR template. qPCR reactions were set up in PowerUp™ SYBR™ Green PCR master mix with gene specific primers (see **Table S2** below), and carried out in AriaMx Real-Time PCR System (Agilent, CA, US). Two technical replicate reactions were carried out for each biological replicate sample. The experiments were done with at least three biological replicates for every treatment group. Each biological replicate comprised of total RNA isolated from cells harvested from one dish.

The RT-qPCR data were analyzed by the ∆Ct method as used earlier (2). In this method, the Ct values that were obtained for CYP1A1 ($Ct^{CYP1A1}$), and the internal control *RPL35a* ($Ct^{RPL35a}$) were processed. Their difference, ΔCt ($Ct^{CYP1A1} - Ct^{RPL35a}$ ) served as a measure of the normalized *CYP1A1* expression in each technical replicate. The average ΔCt of each of the technical replicate provided the normalized expression in each biological replicate. Higher ΔCt value reflected lower *CYP1A1* expression and vice-versa. The ΔCt data obtained for each of the treatment groups were analyzed by one-way or two-way ANOVA, depending on the hypothesis and experimental design.

**Table S2.** List of primers

| Gene | Application | Primer sequence (5´→3´) | Amplicon (base pair) | Annealing temperature (°C) |
|---|---|---|---|---|
| *RPL35a* | RT-qPCR | Forward- CGGCCTCCAAGCTCTCTAAG<br>Reverse- CAGGTCCAGGGGCTTGTACT | 131 | 60 |
| *CYP1A1* | RT-qPCR | Forward-ACCTTTGAGAAGGGCCACATCCG<br>Reverse-<br>TGACTGTGTCAAACCCAGCTCCAAAG | 154 | 60 |
| *CYP1A1* (Amplicon I) | ChIP | Forward- AGTCCCAATTCCAAGGCGTC<br>Reverse- CCTTCGCCATCCATTCCGAT | 406 | 60 |
| *CYP1A1*@ (Amplicon II) | ChIP | Forward- CGTACAAGCCCGCCTATAAA<br>Reverse- CTGGGATCACAAGGATCAGG | 92 | 60 |

@This primer pair was used by Marques et al (3)

#### Total protein isolation and western blotting

The organic (phenolic) phase that is separated out during the isolation of total RNA was preserved for analysis of protein expression. Total protein was isolated from the organic phase as described by Likhite et al (4). Briefly, DNA was precipitated with absolute ethanol, and sedimented via centrifugation. Subsequently, the protein was precipitated by adding isopropanol to the supernatant, and incubation at -20° C for 30 min. The resultant protein pellets were washed with absolute ethanol containing 0.3 M guanidinium chloride. The protein was sonicated and dissolved in 1% SDS. Total protein obtained from each sample was quantified by Lowry's method (5). 30 µg of total protein samples were resolved by 10% denaturing SDS-PAGE, and transferred onto nitrocellulose membrane. The blots were blocked in 1% (w/v) gelatin in 1X Tris-buffered saline containing 0.05% Tween 20 for 2 h at room temperature. Blots were then probed with anti-CYP1A1 antibody, anti-ERα antibody, or anti-AHR antibody overnight at 4 ºC, or for 1h with anti-H3 antibody at room temperature. Blots were washed with 1X TBST (6 washes of 5 min each). Blots were then incubated with HRP-conjugated anti-rabbit, or anti-mouse secondary antibody for 1 h at room temperature. Blots were washed for 30 min (six washes of 5 min each) with 1X TBST, and developed by Clarity Western ECL Substrate.

#### ChIP-seq analysis

ChIP-seq data corresponding to MCF-7 cells treated with E2 (ID: ERR022026), or vehicle (ID: ERR022025) retrieved from Sequence Read Archive (SRA accession ID: ERP000380). The data were analyzed using GALAXY (6). Read quality was assessed using FASTQC (7). Thereafter, the quality scores were converted to Sanger quality type by FASTQ Groomer (8), followed by mapping the reads to reference human genome (hg19) using "Map with Bowtie for Illumina" tool (9). Unmapped reads were filtered out by "Filter SAM or BAM, output SAM or BAM" tool (10). Genomic regions with enriched sequencing reads were identified by MACS (Model-based analysis of ChIP-Seq) tool (11). Resultant Wig files were converted to bigWig files using "Wig/BedGraph-to-bigWig" tool and the peaks representing ERα occupancy were visualized using UCSC genome browser (12).

#### Chromatin immunoprecipitation

Cells were fixed with 0.75% (v/v) formaldehyde for 10 min, followed by addition of 125 mM glycine to stop the reaction. Subsequently, cells were washed with ice-cold DPBS, then lysed with lysis buffer (50 mM HEPES pH 7.5, 140 mM NaCl, 1 mM EDTA pH 8, 1% Triton X-100, 0.1% sodium deoxycholate, 0.1% SDS, 1X protease inhibitor cocktail), and sonicated at an amplitude of 30% for 45 cycles, each cycle with a 10-sec pulse on, and a 25-sec pulse off. Lysates were clarified by centrifugation, and supernatants containing chromatin were collected. 80 µg of chromatin sonicated in ChIP lysis buffer was precleared by incubating with BSA- and herring sperm-coated Protein G plus-Agarose beads. 5% of the pre-cleared chromatin samples were kept aside as input, and the remaining were incubated with anti-ERα, anti-AHR, or normal rabbit IgG antibody for 4 h, at 4˚C. Immune complexes were pelleted by incubating with 20 µl of pre-coated ProteinG plus-Agarose beads for 2 h, at 4˚C, followed by centrifugation. The pellets were washed extensively with a series of wash buffers (13). Immunoprecipitated chromatin samples were eluted and reverse cross-linked as described earlier (13). They were purified using Nucleospin Gel and PCR clean up kit from Machery-Nagel (Duren, Germany). ERα or AHR occupancy was assessed by PCR using two sets of primers relevant to CYP1A1 promoter region (**Table S2**). The PCR reactions were carried out in Veriti 96 Well Thermal Cycler (Applied Bio Systems, USA). The products were resolved on 2% agarose gels. The images of ethidium bromide stained bands were captured using ChemiDoc™ XRS + System with Image Lab™ Software (Bio-Rad Laboratories, Hercules, CA, USA).

#### Statistical analysis

qPCR data with multiple groups, such as those obtained from dose-response experiments were analyzed by one-way ANOVA. However, two-way ANOVA was used to examine the interaction between two variables (EL and 4OHT, EL and fulvestrant, or time and EL). Before applying one-way or two-way ANOVA, the homogeneity of variance was confirmed using the Levene test. TukeyHSD was used for multible comparison. All statistical analyses were performed at 5% level of significance ($p < 0.05$).
