## Supplementary material for "The CYP1A1 connection between enterolactone and breast cancer risk": SI Appendix II

### SI Appendix II: Supplementary figures

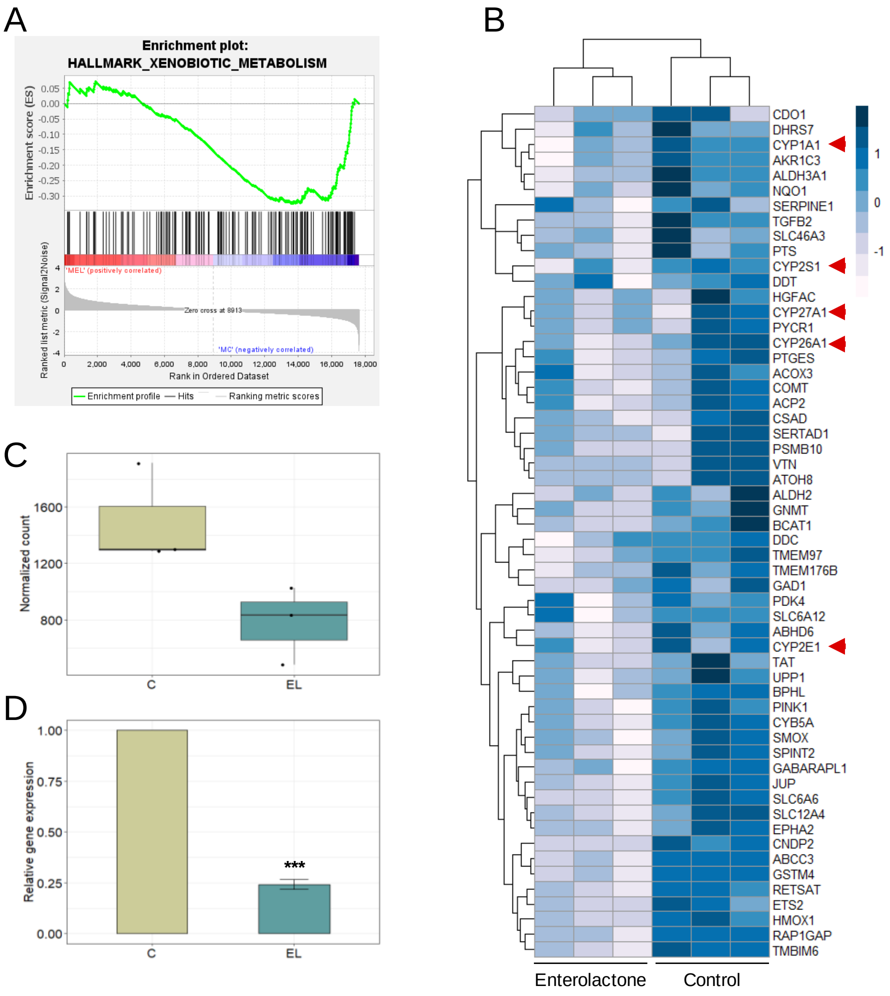


**Fig. S1**: EL modulates xenobiotic response genes in MCF-7 breast cancer cells. In a previous study (1), genes modulated by 10 µM EL in MCF-7 cells were examined for enriched gene-sets using the GSEA software. **A**. Enrichment plot for hallmark xenobiotic metabolism gene-set. **B**. Heatmap showing the pattern of expression of the leading edge genes within the xenobiotic metabolism gene-set in control and EL-treated cells (n = 3 for each). The color bar represents levels of expression. Cytochrome P450 family members are marked by red arrowheads. **C**. A box plot showing the distribution of normalized counts corresponding to *CYP1A1* in three samples each of control and 10 µM EL-treated cells. The mean normalized counts in control- and EL-treated groups are significantly different as inferred from the Wald's t-test, which was as a part of the DESeq2 analysis (padj = 0.01). **D**. RT-qPCR validation. Total RNA samples extracted from control or 10 µM EL-treated cells were subjected to RT-qPCR analysis of *CYP1A1* mRNA expression as described in Materials and Methods (SI Appendix I). The data were analyzed by the relative quantitation $2^{-\Delta\Delta Ct}$ method (2) using *RPL35a* as the internal control. The expression in control cells was set to 1, and that in EL-treated cells were expressed relative to control. Bars represent mean relative *CYP1A1* mRNA expression ± SD (n = 3). The data were analyzed by a one-tailed one-sample t-test to examine whether the mean relative expression in EL-treated samples is significantly less than one. ***p < 0.001. The x-axis labels C, and EL in panels B, C, and D denote control and 10 µM EL-treated cells, respectively.

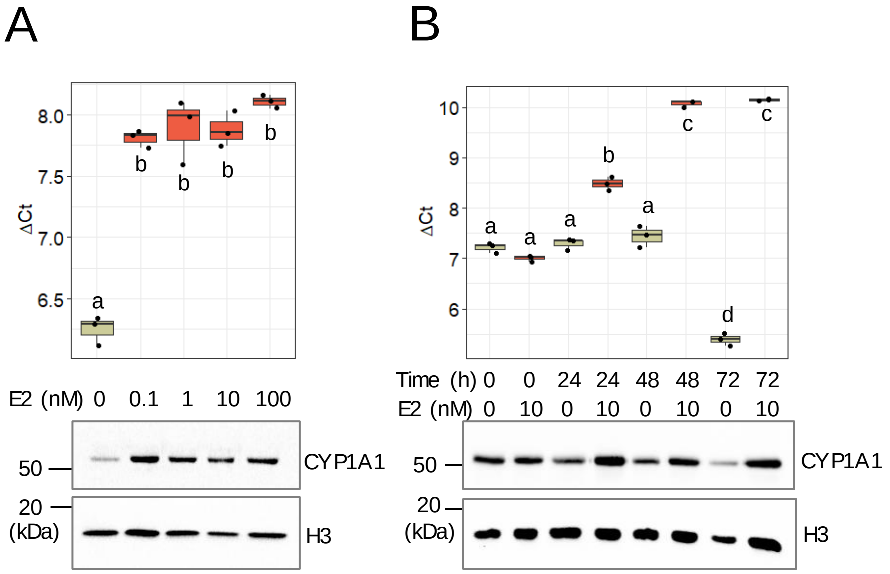


**Fig. S2**. E2 differentially modulates *CYP1A1* mRNA and protein in MCF-7 cells. **A**. Dose-response study. MCF-7 cells were treated with indicated concentrations of E2 in M2 medium for 24 h. **B**. Time-course study. MCF-7 cells were treated with 0.1% DMSO (vehicle) or 10 nM E2 in M2 medium for indicated periods of time. Upon completion of the experiments total RNA and protein were isolated and subjected to RT-qPCR (top panels), and western blotting analysis (bottom panel) to analyze the expression of *CYP1A1* mRNA and protein, respectively. The RT-qPCR data were analyzed by ΔCt method used earlier (1). Here, ΔCt is a measure of *CYP1A1* mRNA expression relative to *RPL35a,* which served as an internal control. Higher the ΔCt value, lower the expression in a given sample. The boxplots show the distribution of ΔCt (n = 3). The RT-qPCR data in **A** was analyzed by one-way ANOVA. The RT-qPCR data in **B** was analyzed by two-way ANOVA to ascertain the main effects of E2, or time, or their interaction. While the main effects of E2 ($p \approx 0$), and time ($p \approx 0$) were significant, their interaction was also significant ($p \approx 0$). Note that in the absence of E2 stimulation, the *CYP1A1* mRNA expression significantly increases with time as reflected by decrease in the ΔCt values. On the other hand, in the presence of E2 stimulation, there is progressive decrease in *CYP1A1* mRNA expression. TukeyHSD was used for multiple comparison. In both the box plots, the statistical difference between pairs of treatment groups are illustrated by lower-case letters. For western blotting, total protein was isolated from the phenolic phase that is separated during total RNA extraction as described in Materials and Methods. The blots were probed with specific antibodies against the indicated proteins. Histone-H3 served an internal control. The chemiluminescence data shown are one of the three biological replicates.

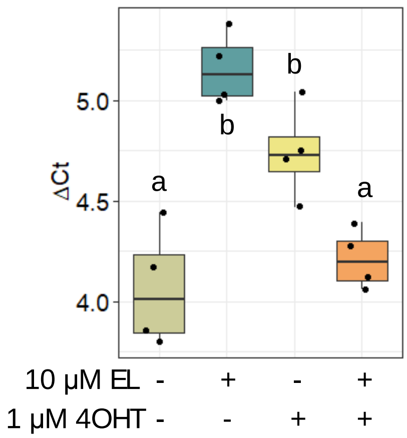


**Fig. S3**. Tamoxifen blocks EL-suppression of *CYP1A1* mRNA. Total RNA was isolated from MCF-7 cells treated with vehicle, 10 μM EL, 1 μM 4OHT, or both for 72 h in M2, and subjected to RT-qPCR analysis. The data were analyzed by ΔCt method. The box plot shows the distribution of ΔCt values ($n = 4$). The data were analyzed by two-way ANOVA to ascertain the main effects of EL, or 4OHT, or their interaction. While there was significant main effect of EL ($p = 0.02$), but not 4OHT ($p = 0.2$), their interaction was significant ($p \approx 0$). a and b denote the statistical difference between pairs of treatment groups, which was determined by TukeyHSD.

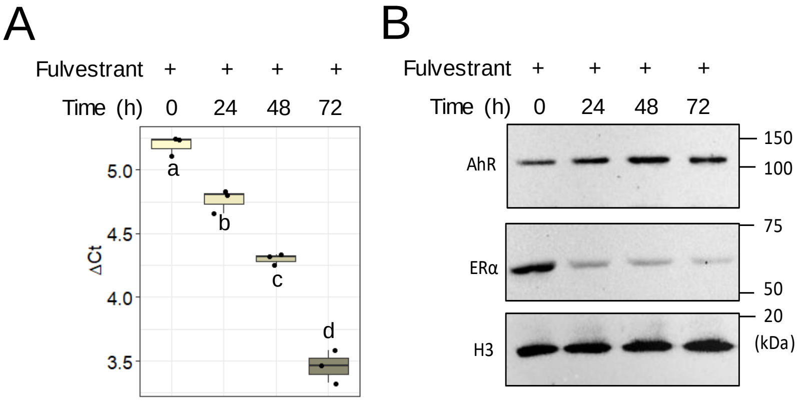


**Fig. S4**: Fulvestrant modulates *CYP1A1* mRNA in MCF-7 cells. Cells were treated with 100 nM fulvestrant in M2 medium for the indicated periods of time. **A**. RT-qPCR. Total RNA was isolated and subjected to RT-qPCR. The data were analyzed by ΔCt method. The boxplot shows the distribution of ΔCt values in each of the treatment groups (n = 3 biological replicates). The data were analyzed by one-way ANOVA followed by TukeyHSD for pair-wise comparison of group means. Letters a, b, c, and d represent statistical difference between pairs of treatment groups. **B**. Western blotting. Total protein isolated from the phenolic phase that is separated during total RNA extraction, was used for western blotting using antibodies specific to ERα, AhR, and Histone-H3 (internal control). The blots were stripped and probed sequentially with AhR and Histone-H3 antibodies. Shown are results obtained from one out of the three biological replicates. Note the increase in AhR protein concomitant to decrease in ERα.

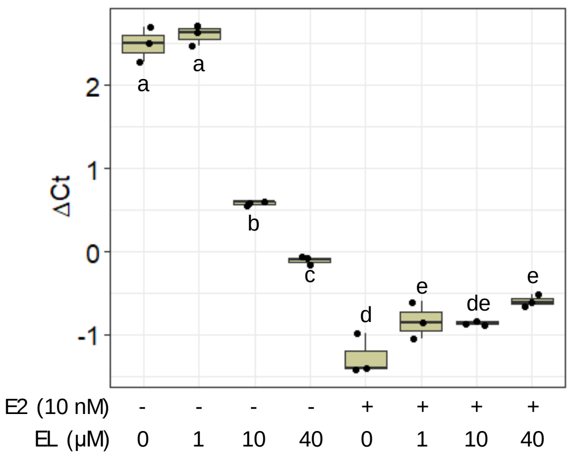


**Fig. S5**. EL reverses the induction of *TFF1* mRNA by E2 in MCF-7 cells. Total RNA was isolated from cells treated with vehicle, or 1,10, and 40 μM EL, alone or in combination with 10 nM E2 for 72 h in M2, and subjected to RT-qPCR analysis. The data were analyzed by ΔCt method. The box plot shows the distribution of ΔCt values (n = 3). The data were analyzed by two-way ANOVA to test the main effects of E2, EL, or their interaction. There were significant main effects of E2 ($p \approx 0$), and EL ($p \approx 0$). The interaction between E2 and EL was also significant ($p \approx 0$). TukeyHSD was used for pair-wise comparison of group means. The lower case letters represent the statistical difference between pairs of treatment. Note that EL, or E2 alone increases the expression of *TFF1* mRNA. However, the increased *TFF1* by E2 is reversed by EL.
